# High tissue-specificity of lncRNAs maximises the prediction of tissue of origin of circulating DNA

**DOI:** 10.1101/2023.01.19.524838

**Authors:** Elyas Mouhou, Fabien Genty, Walid El M’selmi, Hanae Chouali, Jean-François Zagury, Sigrid Le Clerc, Charlotte Proudhon, Josselin Noirel

**Author notes:** These authors contributed equally to this work (Fabien Genty, Walid El M’selmi).

## Abstract

Several studies have made it possible to envision a translational application of plasma DNA sequencing in cancer diagnosis and monitoring. However, the extremely low concentration of circulating tumour DNA (ctDNA) fragments among the total cell-free DNA (cfDNA) remains a formidable challenge to overcome and statistical models have yet to be improved enough to become of practical use. In this study, we set about appraising the predictive value of a variety of binary classification models based on cfDNA sequencing using fragmentation features extracted around transcription start sites (TSSs). We investigated (1) features summarising mapped fragment density around each TSS, (2) long non-coding RNA (lncRNA) genes versus coding genes and (3) selection criteria to generate gene classes to be assigned by the model. Given that, in healthy samples, most of the cfDNA comes from lymphomyeloid lineages, we could identify the model parametrisation with the best accuracy in those lineages using publicly available datasets of healthy patients’ cfDNA. Our results show that (1) the way tissue-specific gene classes are defined matters more than what fragmentation features are included, and (2) in particular, lncRNAs are more tissue specific than coding genes and stand out in terms of both sensitivity and specificity in our results.

**Author summary:** Dying cells, even in healthy individuals, release a fraction of the digested fragments of their genetic material into the bloodstream. Interestingly, these circulating cell-free DNA (cfDNA) fragments bear the footprint left by nucleosomes ; the position of which depends on the transcriptional state specific to each tissue. This footprint, given away by the sizes and genomic positions of cfDNA fragments, can be revealed by deep sequencing and statistical models can be trained to recognise the tissue from where those fragments originate. This information, if made sensitive enough, could be a useful medical technique to carry out so-called “liquid biopsies”, allowing clinicians to diagnose at an early stage or to precisely monitor a number of diseases, including cancer.

In this work, we comprehensively evaluated the features of circulating DNA fragments in the vicinity of transcriptional start sites to increase the ability of statistical models to recognise the tissue of origin of cfDNA of healthy individuals. Broadly speaking, nucleosome patterns allow a statistical model distinguish active versus inactive genes or tissue-specific versus housekeeping genes. The purpose of this study was to find the classes of genes with the strongest ability to recognise the true tissue of origin. From this work, we conclude that long non-coding RNA genes allow for a more sensitive and specific detection of the tissue of origin.

## Introduction

The emergence of precision oncology, which is based on the molecular profiling of the tumours to select adapted treatment interventions, holds great promises to improve cancer treatment. Non-invasive “liquid biopsies”, derived from bodily fluids, allow for the detection of tumour material such as circulating tumour DNA (ctDNA) [1], that is DNA released in the bloodstream by tumoral cells from virtually any tumour site [2]. This technique represents an attractive complement or alternative to invasive and costly tissue biopsies, enabling low-risk tumour profiling from blood and real-time disease monitoring. This opens up the possibility of an optimal, individualised management of patients.

Extensive research has shown the translational potential of ctDNA [3, 4], which is a promising diagnostic and prognostic tool enabling the detection of tumour masses not clinically perceptible [5, 6]. Moreover, longitudinal analyses allow clinicians to follow the evolution of the disease over time. It is possible to assess the early response to treatment, monitor metastatic relapse, or detect the acquisition of resistant mutations [3, 4]. Over the past ten years the most commonly used technologies to detect ctDNA were based on the detection of genetic alterations, which in fact affect only a minor fraction of tumour-derived cell-free DNA (cfDNA). In particular, epigenetic alterations, such as DNA methylation changes, or fragmentation patterns provide an additional level of information [7]. Epigenetic landscapes are also highly cell-type specific and inform on the tissue of origin [8–10]

A growing body of work has shown that cfDNA fragmentation patterns can be used to infer what cell-type, tissue or organ is likely to have released these fragments in the bloodstream. This can help deduce whether someone is healthy or not and locate the tumour when someone has cancer. Indeed, there exists a broad statistical association between the expression level of gene and how the nucleosome particles are organised around its transcription starting site (TSS) [11, 12]. As it happens, the majority of cfDNA derives from dying cells through apoptosis and its fragmentation patterns thus inform us about nucleosome positioning in those cells [13]. As a consequence, plasma DNA sequencing data can in principle be harnessed to sketch an expression landscape for the cells that have released cfDNA into the bloodstream. This expression landscape can in turn be used to identify the likely tissue or tissues of origin of cfDNA. In previous studies, cfDNA sequencing data was used to characterise nucleosome positions, nucleosome spacing and depth of sequencing around the TSS of coding genes [14, 15]. In fact, it is not just the genomic regions surrounding TSSs that can used but all regulating regions bound by transcription factors [16]. Finally, larger-scale, position-dependent fragmentation features can also carry information as to the cancer status of a patient [17].

The ENCODE project established that the majority of the genome is functionally active even if not to code for proteins [18]; this led to a concerted effort aiming at characterising functional and regulatory units in non-coding DNA [19]. Among those, many loci encode long non-coding RNAs (lncRNAs), RNA transcripts more than 200 nucleotides long which do not encode any protein. An association with lncRNAs has already been been found for virtually every single disease [20] and their expression (albeit lower than that of coding genes) appears to be highly tissue specific [21, 22].

Annotations of lncRNAs are provided in several databases [19]. FANTOM CAT compiles an atlas of more than 25,000 lncRNAs with high-confidence 5^*1*^ ends and expression profiles across samples from many primary cell types and tissues [23].

As a follow-up to previous work aimed at recognising the tissue of origin of cfDNA based on fragmentation patterns around TSSs [14, 15], we addressed here the question of how tissue specificity could be used and defined to maximise the prediction accuracy. To this end, we comprehensively studied the characteristics of circulating DNA fragments around TSSs that increase the ability of statistical models to recognise the tissue of origin of cfDNA in healthy individuals. We evaluated how the sensitivity of the method depended on what sets of genes were to be classified. Besides and more importantly we investigated the value of using the tissue-specificity of lncRNAs.

## 1 Materials and methods

### 1.1 Overview

We used Plasma-Seq data derived from blood samples of healthy individuals previously produced by Snyder et al. [15]. Fragmentomic profiles were computed including a fragmentation protection score (as described in [15]), reflecting patterns of fragmentation of cell-free DNA. Those profiles, restricted to ± 5 kbp around transcription start sites (TSSs) of coding and non-coding genes, were computed by integrating information from FANTOM [23, 24]. Each gene’s fragmentomic profile was processed to provide a characterisation based on a dozen of fragmentomic features (see Fig. 1 and section “Fragmentomic features” below).FANTOM provided us with expression data corresponding to the major human primary cell types and tissues, known as “ontologies” in FANTOM parlance. FANTOM also provided information regarding the tissue specificity of each gene, which allowed us to define, together with the gene category (coding or non-coding), for each ontology, a number of tissue-specific classes of genes (see section “Gene classes based on transcription and tissue specificity” below).

**Fig 1.**
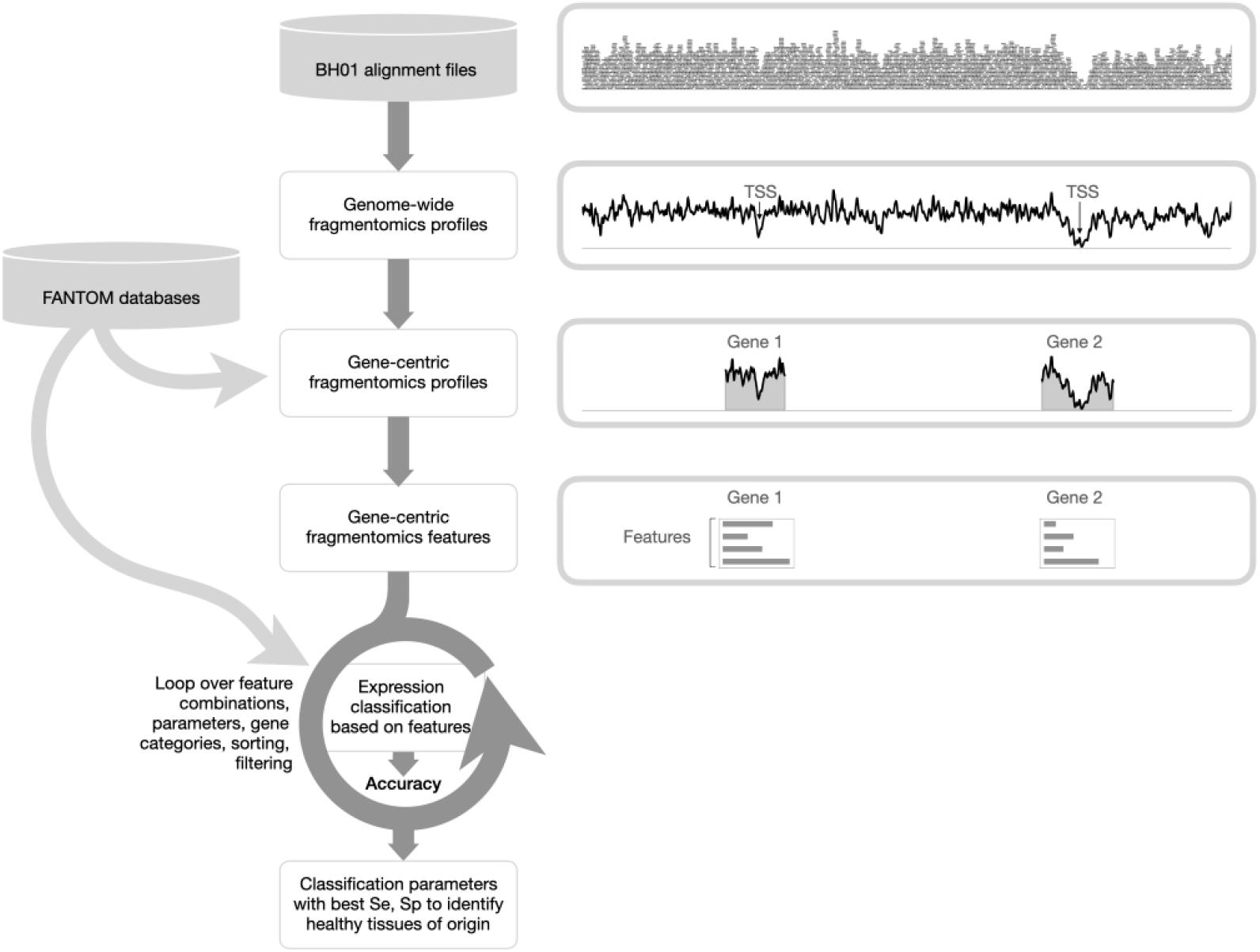
Overview of the pipeline. First we processed the Plasma-Seq data into gene-centric fragmentomic features. Those features were used for each FANTOM ontology to classify gene classes specific of said ontology. In order to measure the performance of a model configuration we set about increasing the ability of the model to accurately classify genes in lymphomyeloid-related ontologies while decreasing its ability to accurately classify genes from unrelated cell types.

A model configuration corresponded to a particular set of features and a particular choice of gene classes. Each model configuration was used to train a support vector machine (SVM) to classify genes in each ontology. The classification accuracy was evaluated by random resampling. The output of our pipeline was a table of accuracy values for each ontology and each model configuration. Our objective here was to identify the best model configuration, ie, the model configuration for which the tissue or tissues of origin had a good accuracy both in absolute terms and relative to other, non-relevant tissues.

### 1.2 Datasets and databases

#### 1.2.1 Plasma-Seq data

In this study, we re-analysed the dataset BH01 obtained and described in [15]: it corresponds to cfDNA extracted from a pool of plasma from several healthy donors, deeply sequenced (100x) using paired-end reads of length 101 nt. Unprocessed read data are available from the NCBI Sequence Read Archive under the accession number SRX1120757 from the PRJNA291063 bioproject. In this study, we started off with the BAM files available from Martin Kircher’s [website](https://kircherlab.bihealth.org/download/cfDNA/) (accessed 2018). Information regarding data processing in the original article can be found on Jay Shendure’s laboratory’s GitHub repository [cfDNA](https://github.com/shendurelab/cfDNA).

#### 1.2.2 The FANTOM databases

The FANTOM Consortium aims at producing a functional annotation of the mammalian genome that extends ENCODE’s own endeavour [18, 25]. The research and the data produced since 2000 by the FANTOM Consortium have been packaged as iterations, from FANTOM1, a functional annotation of a mouse cDNA collection, to FANTOM6, a functional analysis of non-coding RNAs. For this study, we relied on the output of FANTOM5, which compiles an atlas of mammalian promoters, enhancers, lncRNAs and miRNAs using a cap analysis gene expression (CAGE) approach [23, 24]. More specifically, FANTOM5 provided us with a list of genes, categorised as coding or non coding in order to evaluate the benefit of using one category or the other in terms of tissue specificity. Additionally, FANTOM5’s CAGE data allowed us to get an idea of the location of each gene’s transcription start site (TSS), and we needed this because the features used for the tissue identification are fragmentomic features extracted around the TSSs. Technically, TSSs were identified from clusters of decomposition-based peaks (DPI for decomposition-based peak identification). Finally, the CAGE expression profiles were available for a number of cell lines and tissues, known as “ontologies”, which allowed us to derive lists of active or inactive genes (referred to as “gene classes”) for each tentative tissue.

TSS positions and gene categories were obtained from processing three data files originating from the FANTOM project [23]: “FANTOM DB1” a database providing for each gene ID the DPI cluster corresponding to the most robust TSS, “FANTOM DB2” a database providing for each gene ID the gene category (coding genes, pseudogenes, etc.) and “FANTOM DB3” a database providing from CAGE alignment the genomic locations of the DPI clusters. All three databases were combined for the purpose of associating each of the 59,110 genes to a gene class (see “Gene classes based on transcription and tissue specificity”) and its TSS’s genomic location. (See “Code and data availability” for URLs.)

The FANTOM5 project [23, 26] has produced an atlas of coding and non-coding RNA using the cap analysis of gene expression (CAGE) technology in a variety of well-defined cell and tissue samples, referred to as “ontologies”, corresponding to 173 Cell Ontology terms [27] and 174 Uberon terms [28]. This allowed Hon et al. to record the cell-type specificity of each genes both in terms of a fold-change and a *p*-value [23]. Briefly, for each ontology, the expression (in units of TPMs, “transcripts per kilobase million”) in the corresponding samples were compared to the expression in the rest of the samples with a Mann-Whitney sum rank test, providing the *p*-value. The fold-change for an ontology was obtained by dividing the mean TPM of the associated samples by the mean TPM of the other samples. In this study, we relied on “FANTOM DB4”, a set of 347 data files recording for each ontology the specificity of each one of the 59,110 genes in terms of fold-change and *p*-value.

We also analysed fragmentation patterns around the TSSs of housekeeping genes. We built a gene class of “ubiquitous genes” using the database of housekeeping promoters “FANTOM DB5” (supplementary table 4 in [24]). This table of 10,787 genes was filtered to keep only the strongest DPI cluster of the robust FANTOM set (our FANTOM DB1) leaving us with 6524 genes, including 5884 coding genes and 53 lincRNAs. The final table contained 5711 coding genes and 48 lincRNAs (when intersected with our using our FANTOM DB2, which only considers the genes located on the autosomes).

Finally, we used expression data to provide a convenient 2D representation of the ontologies, onto which we mapped some of our key results. The data tables “FANTOM DB6a” (count data in TPM for each sample) and “FANTOM DB6b” (sample-ontology relationship) provided by [23] were used for that purpose. See the Data and code availability section.

#### 1.3 Fragmentomic profiles

The end game of this study was the identification of the tissue of origin of cfDNA. Since the level of expression of a gene, which is tissue-specific, is associated with a more or less precise nucleosome positioning around this gene’s TSS and since this positioning in turns affects how the genome is fragmented upon apoptosis, we sought to capture susceptibility to fragmentation around each gene’s TSS from the sequence data.

The sequences of cfDNA fragments were used to calculate genome-wide fragmentomic profiles from the DNA reads mapped in proper pairs corresponding to inserts of sizes comprised between 120 nt and 180 nt of BH01. This included (1) the “fragment depth” (FD), defined at each genomic position as the number of inserts that overlap this position [14] and (2) the “window protection score” (WPS) defined in [15], which measured for each genomic position the susceptibility to fragmentation of the region centred around it. More precisely, given a window size *w* at each genomic position, the WPS was calculated as S = *O* − *F*, where *O* denotes the number of inserts that fully overlapped the window and *F* the number of inserts that only partially overlapped with the region (only one end fell within the region). A width *w* = 101 was chosen, in accordance with Snyder et al. [15].

Fragmentomic profiles were then restricted to regions centred on the TSS of each coding gene or intergenic lncRNA. This was done using the FANTOM databases DB1 and DB3, assuming the TSS corresponded to the position of the strongest DPI (decomposition based peak identification) cluster identified in Ref. [23]. Besides, for merely technical reasons, the analysis pipeline at this stage operated separately on the 21,069 coding genes and on the 13,105 intergenic lncRNAs using the gene category provided by FANTOM DB2. TSS-centred regions were taken to be 10,001 nt wide: 5000 nt upstream and 5000 nt downstream of the TSS, in addition to the TSS itself. The protection score was calculated over these regions, in accordance to what had been described in a previous study [15]. (Nucleotides were counted relative to the TSS with position zero corresponding to the first transcribed nucleotide.) Genes located on the reverse strand of each chromosome were turned around before proceeding with data processing.

#### 1.4 Fragmentomic features

From the fragmentomic profiles we derived twelve “fragmentomic features” on a small, medium and broader scale for each gene. More precisely, four regions were identified on which summary statistics were computed (see Fig. 2: a “distal region” corresponding to 10,000 nucleotides around the TSS, a “proximal region” corresponding to 2000 nucleotides around the TSS, a “TSS region” from 150 nucleotides upstream to 50 nucleotides downstream of the TSS and a “*n* + 1 region” (corresponding to the region were the first nucleosome would get evicted) from the TSS to 300 nucleotides downstream of it.

**Fig 2.**
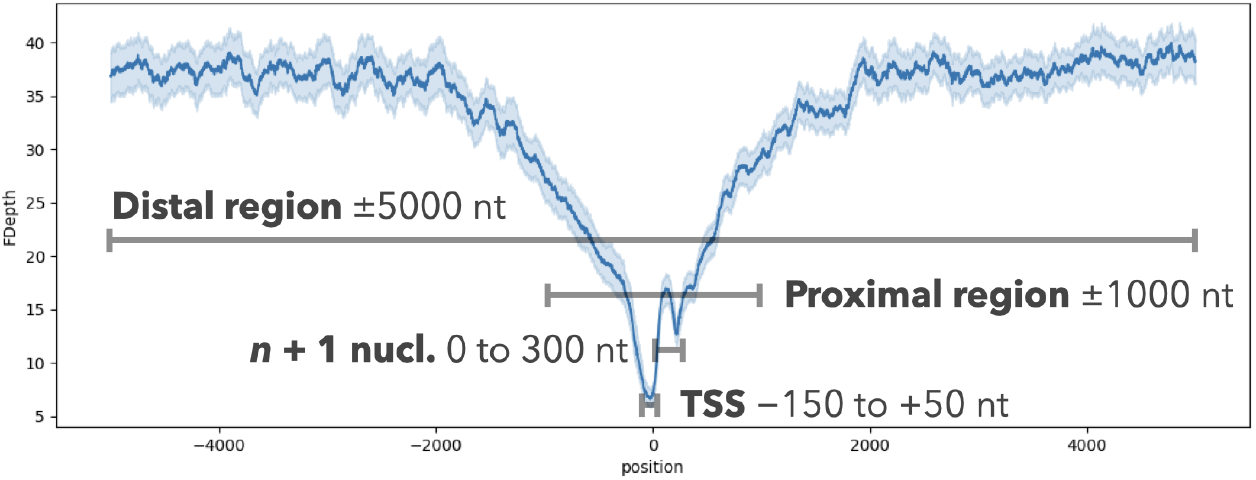
Canonical profile in the nucleosome depleted region around a TSS. Summary statistics are computed at various scales around the TSS to characterise fragmentation patterns for a given gene. Those statistics are then used as fragmentomic features.

The first three features measured the intensity of fragmentation around the TSS:

- 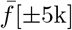, the FD averaged over the region that runs from − 5000 to +5000 nt relative to the gene’s TSS,
- 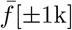, the FD averaged over the − 1000 to +1000 nt region,
- 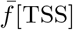, the FD averaged in the vicinity of the TSS (−150 to +50 nt region). The next two features measured the dispersion of the FD around the TSS:
- *s*^2^[TSS], the variance of FD in the vicinity of the TSS,
- *r*[TSS], the range of FD in the vicinity of the TSS, Scaled features were derived from the last four features described above by normalising them using the broader-scale average 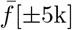:
- 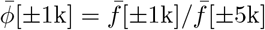, the average normalised FD in the ±1k region,
- 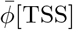, the average normalised FD in vinicity of the TSS,
- *σ*^2^[TSS], the variance of the normalised FD *φ* in the vicinity of the TSS and
- *ρ*[TSS], the range of the normalised FD *φ* in the vicinity of the TSS. We defined the average FD in the vicinity of the TSS relative to the average computed over the ±2k region:
- 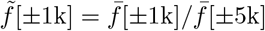, the ratio of FD averages in the ±1k region.

Two features were computed using the WPS:

- ∆[0→ +300], the range of WPS in the 0 to +300 nt region (corresponding to the position of the first nucleosome past the TSS) calculated as max *W*_*i*_ − min *W*_*i*_ where the *W*_*i*_ denotes the WPS at position *i* and where the maximum is taken over the 0 to +200 nt region relative to the TSS and the minimum is taken over the +150 to +300 nt region relative to the TSS.
- *δ*[0 → +300], a similar quantify but calculated from WPS values normalised by the average WPS over the ±5k region.

#### 1.5 Fragmentomic feature sets

Fragmentomic features were combined in twelve “feature sets” (FSs) to identify the most predictive ones. The list of FSs is given below and Table 1 provides a tabulated summary.

**Table 1.**
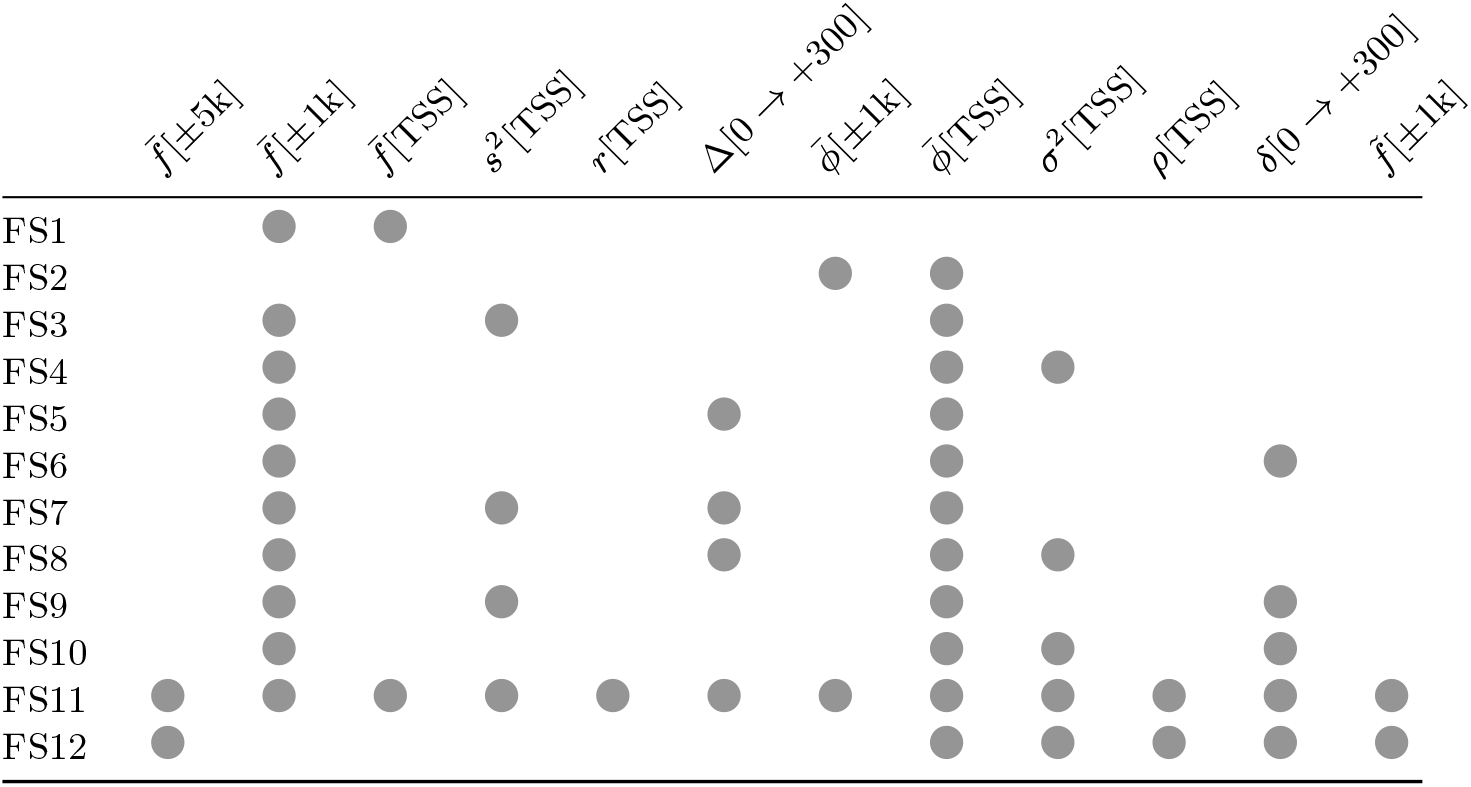
Description of the feature sets analysed in this study.

FS1 with 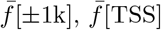

FS2 with 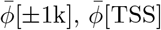

FS3 with 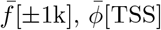, *s*^2^[TSS]

FS4 with 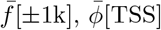, *σ*^2^[TSS]

FS5 with 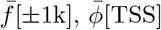, ∆[0 → +300]

FS6 with 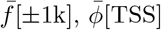, *δ*[0 → +300]

FS7 with 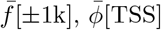, *s*^2^[TSS], ∆[0 → +300]

FS8 with 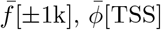, *σ*^2^[TSS], ∆[0 → +300]

FS9 with 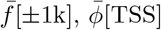, *s*^2^[TSS], *δ*[0 → +300]

FS10 with 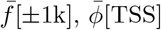, *σ*^2^[TSS], *δ*[0 → +300]

FS11 with 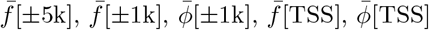, *s*^2^[TSS], *σ*^2^[TSS], *r*[TSS], *ρ*[TSS], ∆[0 → +300], *δ*[0 → +300], 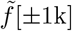 (all features are included)

FS12 with 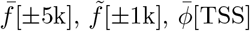, *σ*^2^[TSS], *ρ*[TSS], *δ*[0 → +300]

#### 1.6 Gene classes based on transcription and tissue specificity

If a cell type contributes significantly to the cell-free circulating DNA, it is expected that genes of similar transcriptional activity will exhibit similar fragmentation patterns in the vicinity of their TSS. This similarity can be picked up, in principle, by learning algorithms trained to effectively classify genes, say, as transcriptionally active or inactive based on fragmentomics features alone.

In this paper, rather than settling for a single denotation of transcriptional activity, we decided to operationally and functionally diversify how gene classes were defined so as to encompass notions of transcriptional activity and more importantly tissue-specificity. Indeed what really mattered for the application at hand was whether fragmentation patterns can be easily distinguished for some classes of genes in the tissue to be identified.

For each single FANTOM ontology (347) used in this study [23], FANTOM DB4 provides a dataset informing us on each gene specificity in terms of fold-change and value. These datasets were used to define a number of “gene classes”, which depend on what ontology selected (see Fig. 3):

**Fig 3.**
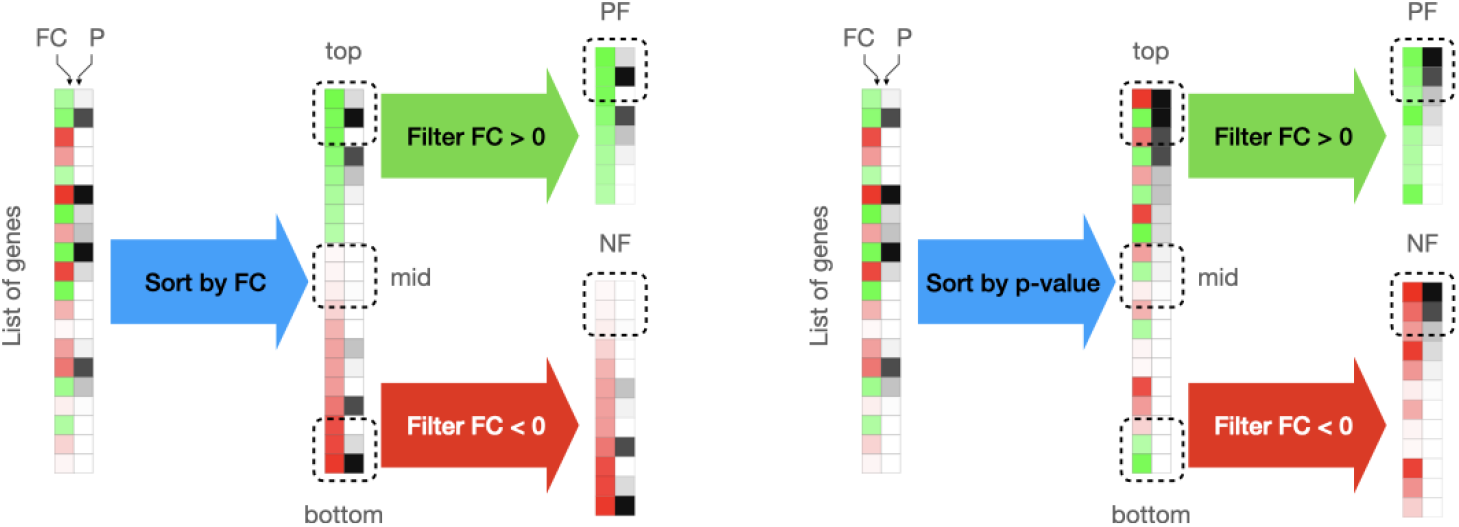
From tissue specificity to gene classes. FANTOM provides for each ontology a list reporting the tissue specificity of each coding gene in terms of fold change (left) and *p*-value (right). In this diagram, the heatmaps correspond to one such list where colour represents the fold-change: the hue increases with the absolute fold-change and colour (green or red) encodes the sign of the fold change while the grey levels represent statistical significance (darker means lower *p*-values). From this list, we defined a number of gene classes (top, mid, bottom, PF and NF) by using the fold change (higher fold changes at the top of the list) or the *p*-value (lower *p*-values at the top of the list) as a sorting criterion optionally filtered based on the sign of the fold change. Each gene class ultimately contains 100 lincRNAs or coding genes, depending on the gene category under consideration.

- Select a gene category: either coding genes (21,069) or intergenic lncRNAs (13,105).
- Sort by decreasing fold-change “FC sorting” (high fold-changes appear at the top) or increasing *p*-value “P sorting” (significant *p*-values appear at the top); this allowed us to define three possible classes: (1) top, (2) bottom or (3) middle of the list.
- Additionally, focus on positive fold changes (genes that are specifically over-expressed in an ontology) or negative fold changes (genes that are specifically under-expressed in an ontology). This allowed us to define two other classes: positive-fold (PF) and (5) negative fold (NF) genes.
- Regardless of the sorting criterion, (6) a list of 100 random ubiquitous, housekeeping genes was extracted from the list provided by [24] as described above. This list was kept identical throughout this study (see Supporting Information S1).

For each category (coding genes or lincRNAs), for each sorting criterion (fold-change or *p*-value), for each ontology, the following gene classifications were trained and evaluated: positive-fold vs bottom (PF:b), negative-fold vs bottom (NF:b), positive-fold vs negative-fold (PF:NF), positive-fold vs ubiquitous (PF:u), negative-fold vs ubiquitous (NF:u), bottom of the list vs ubiquitous (b:u), positive-fold vs middle of the list (PF:mid), negative-fold vs middle of the list (NF:mid) and bottom vs middle of the list (b:mid).

#### 1.7 Gene classification

For each ontology and each model configuration, classification was performed using a support vector machine (SVM) with a non-linear radial basis function as kernel and a regularisation parameter *C* = 1, as implemented in Scikit Learn. The classifier’s performance was measured in terms of accuracy, which is evaluated through random permutation cross-validation with 100 splits, each one constituted of 70% training and 30% test data.

#### 1.8 Indicators for model selection

Among the 347 ontologies used in this work, we selected 40 belonging broadly to the lymphomyeloid category which we targetted [29]; eg, hematopoietic stem cell (CL 0000037), common myeloid progenitor (CL 0000049), leukocyte (CL 0000738) or blood (UBERON 0000178). This set of ontologies to the identifiable “lymphomyeloid cluster” in the *t*-SNE map described below (see also Fig. 4) and is provided as Supporting Information S1.

**Fig 4.**
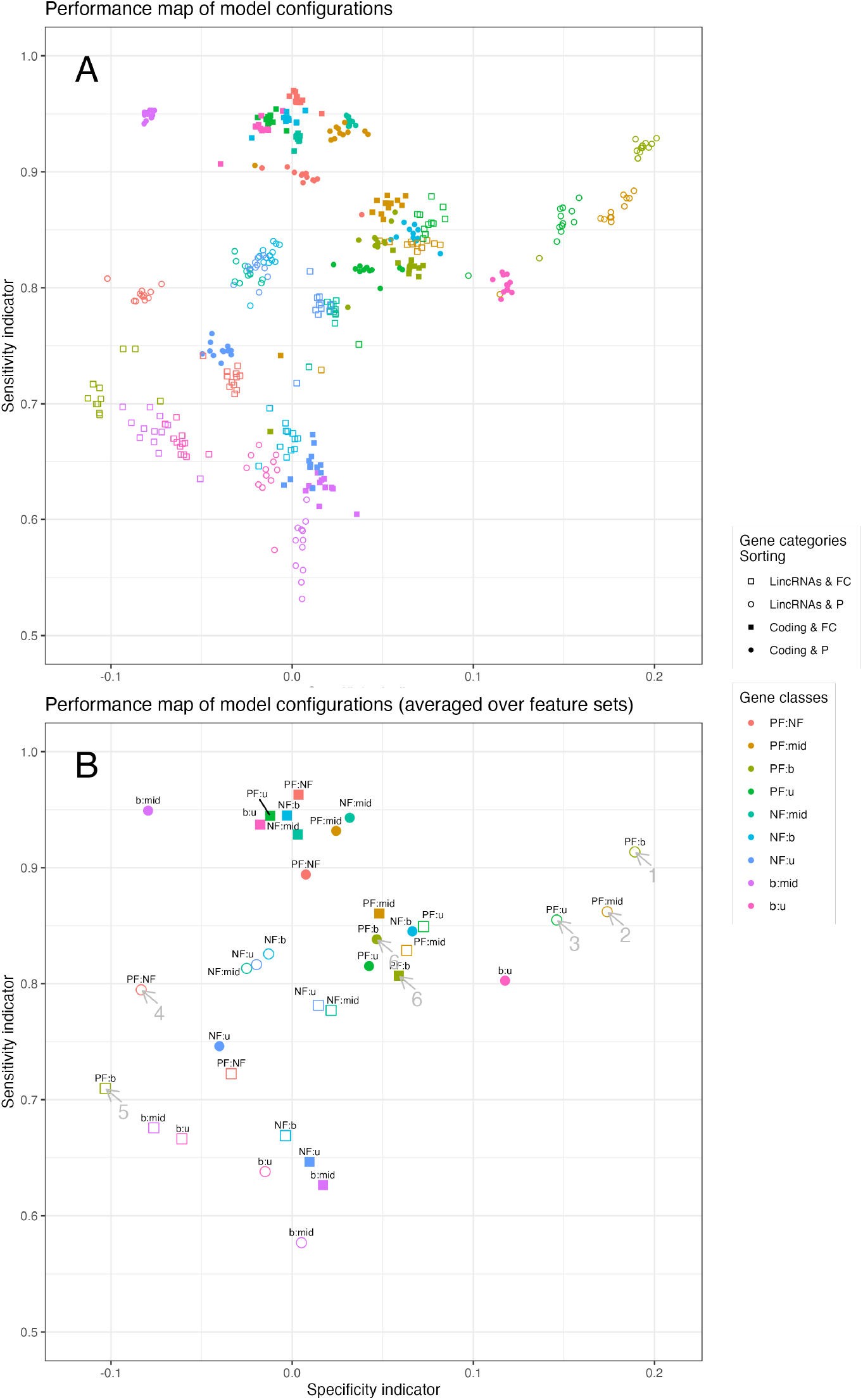
Performance map of model configurations. The sensitivity and specificity indicators were used to display the efficiency of each model configuration. The sensitivity indicator quantifies how good gene classification can get in target ontologies belonging to the lymphomyeloid cluster whereas the specificity indicator quantifies the gap between target ontologies and non-target ontologies. The best model configurations will therefore appear in the upper right corner of the map. (A) Sensitivity and specificity indicators for all model configurations. Shapes were used to distinguish gene categories: lincRNAs use hollow symbols, coding genes use filled symbols; sorting by fold-change uses squares, sorting by *p*-value uses circles/discs. Colours were used to distinguish the pairs of gene classes the model is trained to predict. For clarity, feature sets were not individualised. (B) Sensitivity and specificity indicators averaged over feature set parameters, since model configurations differing only in the feature sets they use tend to cluster together. Arrows indicate model configurations discussed more at length in the text. Our GitHub repository includes a Bokeh rendition of the results that allows the user to dynamically interact with the graph.

A suitable model configuration should yield a high accuracy for lymphomyeloid ontologies and a low accuracy for the other ontologies, since the sequencing data were been collected from healthy individuals. We therefore computed two indicators that play a role loosely reminiscent to that of sensitivity and specificity and, in the rest of this paper, we use the terms “sensitivity indicator” and “specificity indicator” to refer to them:

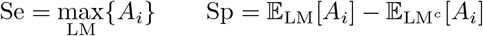

where expectation and maximum are calculated over the lymphomyeloid cluster (LM) — the set of 40 ontologies defined above — or over its complement (LM^*c*^), ie, other ontologies. We have, on the one hand, the expected accuracy of the method for the target ontologies and, on the other hand, the expected gap in accuracy between the target ontologies and other ontologies. The model configuration we wished to retain is one with high sensitivity and high specificity, so that the target ontology can be identified with a comfortable margin compared to irrelevant ontologies.

#### 1.9 Graphical map of model performance

In order to represent in a visually meaningful way the performance of the best model configuration to classify gene classes, we used a *t*-stochastic neighbour embedding (*t*-SNE) of FANTOM’s expression data to derive a 2D map of the FANTOM ontologies, on which we could plot model accuracy estimates. This map was also useful to identify the cluster of lymphomyeloid-like ontologies. For each gene and for each ontology, we calculated the median expression across the samples associated with said ontology (expression levels are expressed in transcripts per million or “TPM”). Genes exhibiting zero variation across the ontologies were removed from the analysis. The *t*-SNE was then calculated from log-transformed median expression values.

## 2 Results

To identify the tissues contributing to the pool of circulating DNA, we aimed at developing a model able to recognise in a sensitive and specific manner the tissue (or tissues) of origin from the fragmentation patterns around TSSs that arise from nucleosome positioning, transcription factor binding or chromatine accessibility.

Recognising this tissue of origin required us to focus on tissue-specific patterns and first and foremost on tissue-specific genes. This study pragmatically explored the question of how best to define “tissue specificity”, how to leverage tissue specific gene classes and how to fine-tune the model’s parameters in order to make headway in terms of sensitivity and specificity indicators. We did this by testing all 428 possible combinations (hereafter referred to as “model configurations”) of feature sets, gene categories and tissue-specific gene classes on data collected from healthy individuals and by charting the performance of each model configuration in terms of sensitivity and specificity indicators, shown in Fig. 4.

### Importance of factors determining performance

With the sensitivity indicator as the abscissa and specificity indicator as the ordinate, the best model configurations were found towards the upper right corner of the plot in Fig. 4. In Fig. 4A, feature sets were not individualised for clarity. It appears that model configurations using the same sorting method, the same gene classes and the same gene categories (ie, differing only in terms of the feature sets they use) cluster tightly together on the performance map.

Among the various factors determining the model configuration, gene categories, gene classes and sorting were by far the primary factors determining performance. Feature sets, which we discuss below, had themselves little impact on performance. It made sense therefore to simplify this map by averaging the performances over the feature sets (Fig. 4B). See Supporting Information S2 and S3 to explore our results.

### Best performing model configurations all use lincRNAs and P-sorting

Discounting all model configurations that lined up along a specificity indicator of zero, we observed that the top three model configurations (upper right corner of the plot) all relied on lincRNAs together with P sorting and one of the gene class being PF (cf. Fig. 3). In those highly sensitive and highly specific model configurations, the PF gene category was contrasted against one of the following gene classes:

- the bottom of the list gene class (b, arrow 1),
- the middle of the list gene class (mid, arrow 2),
- the list of ubiquitous genes (u, arrow 3).

The fourth possible gene class would naturally be PF:NF and, while this would have seemed a natural candidate to bet on, its performance was surprisingly poor (with Sp *<* 0, arrow 4).The observation that best model configurations all depended on lincRNA genes can be largely explained by the high-tissue specificity of lincRNAs [21]. However, it should be noted that tissue specificity isn’t sufficient in itself to secure good performance.

Indeed, model configurations using an identical parametrisation except that they used sort by fold-change instead of *p*-value all displayed poor performance (eg, the PF:b classification indicated by arrow 5). The performance of PF:b classification using lincRNAs thus critically depends on sorting. This must be contrasted to the PF:b classification problems using coding genes, which were consistent whether one chose to process the lists using FC-sorting or P-sorting (arrows 6). It remains to be seen why P sorting appears superior to FC sorting specifically for lincRNAs. One way to approach this question was to go back to the average fragmentation profile for each class of gene in an ontology representative of the lymphomyeloid cluster.

### Classification performances and average fragmentation profiles

The efficacy of a model configuration depended substantially on how much, in any relevant target ontology, the typical fragmentomic profile observed among genes of one gene class will differ from that observed among genes of the opposing gene class. In order to test this, we computed and compared the average profiles of each gene class in the neutrophil ontology Fig. 5.

**Fig 5.**
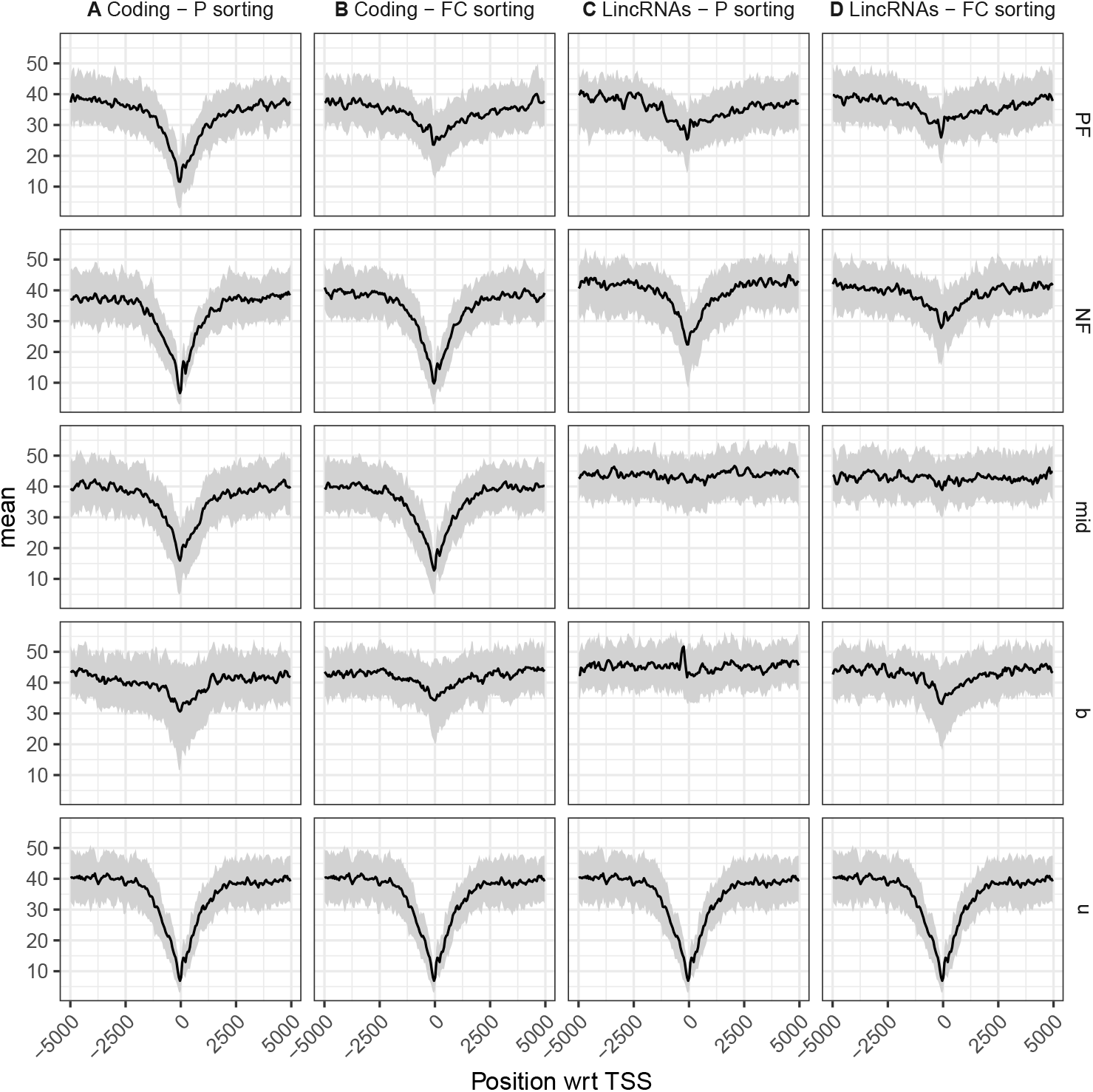
Fragmentomic profiles for the neutrophil ontology. Average fragmentomic profiles (and its 95% envelope) around the TSS for each of the gene classes PF, NF, b, mid and u for the neutrophil ontology (CL 0000775), among coding genes or lincRNAs, sorting by fold-change or by *p*-value (columns A, B, C, D). The fragment depth was used for that purpose and position is relative to the TSS genomic position (see Materials and methods). Note that the ubiquitous “u” category is identical regardless of gene category and of sorting criterion, so the profile is identically repeated four times.

Generally speaking, the fragmentomic profiles either looked flat or displayed a more or less pronounced depression in terms of fragment depth (FD).

One of the striking differences in the profiles could explain the superiority of lincRNAs over coding genes in the PF:b classification. We observed that for lincRNAs sorted by *p*-value, the profiles of “b” genes was flat in contrast to “PF” genes, which displayed a distinctive depression in the vicinity of the TSS (PF:b in Fig. 5C). This could explain the high accuracy for this model configuration using lincRNAs (arrow 1 in Fig. 4). It must be noted that the profiles of “b” genes weren’t flat when sorting by fold-change, which resulted in deteriorated accuracy (PF:b in Fig. 5D, see arrow 5 in Fig. 4). For coding genes, a different picture emerged. The average fragmentomic profiles were similar for both gene classes, both displaying a depression around TSSs. Importantly this was the case regardless of the sorting method (PF:b in Fig. 5A and B). This would explain why the PF:b classification behaved consistently in both model configurations (arrows 6 in Fig. 4).

The performance map could leave the reader under the impression that the middle of the list and the ubiquitous (mid and u) classes are similar in terms of fragmentomic signatures. Indeed, many of the pairs of model configurations obtained by simply substituting the mid class for the u class tended to cluster together on the performance map. See, for instance, the following pairs in Fig. 4B: PF:u and PF:mid in P-sorted lincRNA classes or NF:u and NF:mid in FC-sorted lincRNA classes). However, the average profiles helped us see that this similarity was only coincidental for lincRNAs. The fragmentomic profile of ubiquitous genes was distinctively prototypical of an actively expressed gene (Fig. 5[u]). In contrast, the fragmentomic profile of middle of the list lincRNAs was breathtakingly flat, regardless of sorting (Fig. 5C[mid] and Fig. 5D[mid]).

### Performance across feature sets

Even though the impact of the specific choice of features used to classify genes appeared marginal, patterns in terms of sensitivity and specificity indicators emerged in this analysis. We computed the changes in sensitivity and specificity indicators attributable to each feature set (averaged over all the other parameters of a model configuration). One feature set, FS2, stood out and performed far worse than the others; FS2 is made only of the two scaled quantities 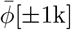 and 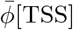(Fig. 6A). FS11 (all features included) and FS12 appear to perform better (Fig. 6B).

**Fig 6.**
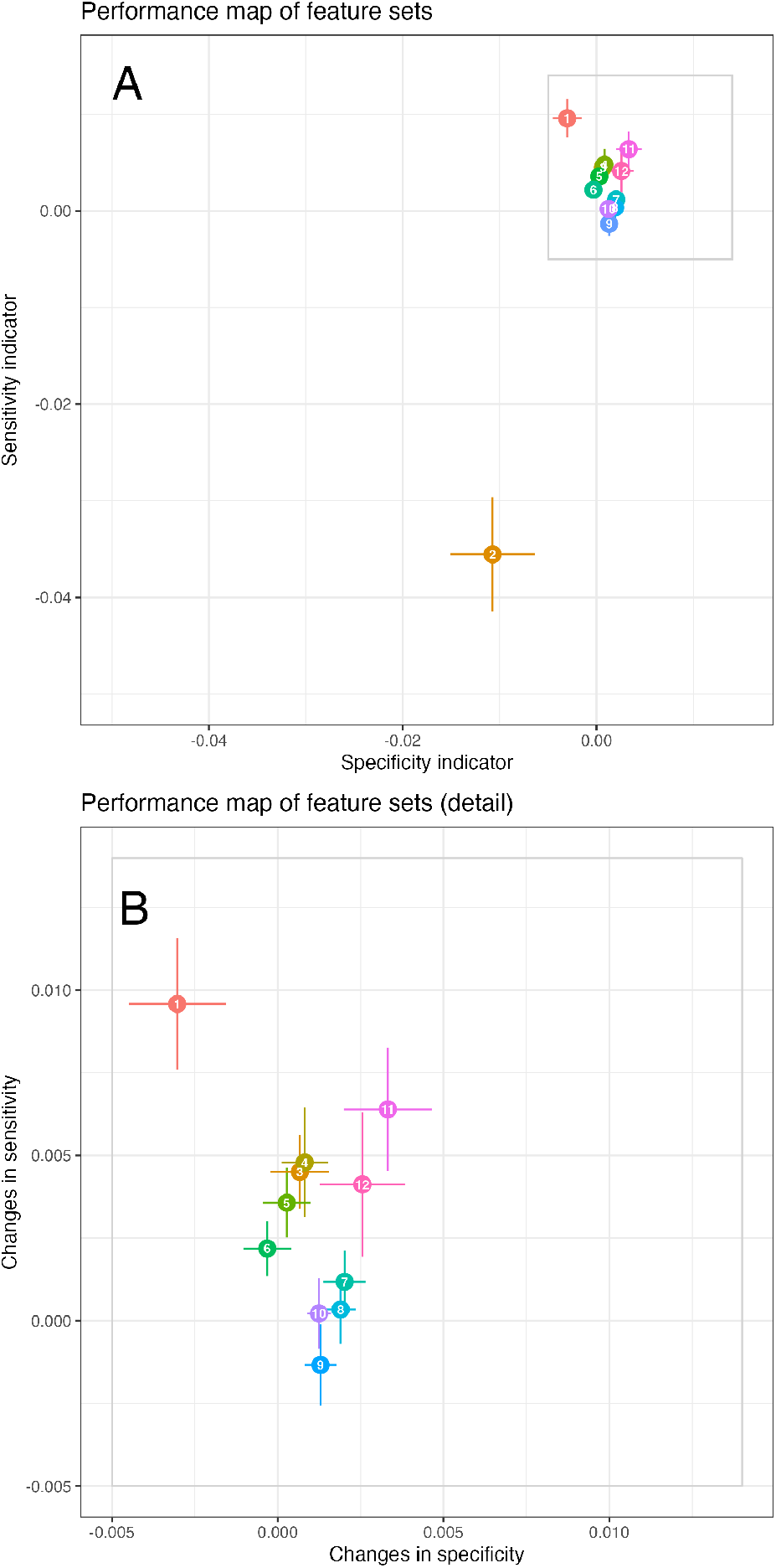
Performance changes attributable to feature sets. (A) For each feature set, performance indicators were centred with respect to all other model configuration parameters, in order to measure changes in performance that could be attributed to each feature set. Error bars correspond to standard errors in sensitivity or specificity. The feature set FS2 was associated with strikingly poor performance estimates and, as a consequence, (B) a detailed version of the previous plot, omitting FS2 is provided.

### Mapping performance across ontologies

The approach described above allowed us to identify the model configuration with the best combination of sensitivity and specificity indicators: lincRNAs, P-sorting and PF:b gene classes. In order to better how good sensitivity and specificity could be achieved by this model configuration, we charted the accuracy for each FANTOM ontology on an 2D chart, see Fig. 7A. (See Supporting Information S4 for an interactive plot.) The coordinates of each ontology was obtained using *t*-SNE as described in the Materials and Methods. On this chart, functionally-related ontologies tend to cluster together. In particular, the brain ontologies and lymphomyeloid samples markedly stand out.

**Fig 7.**
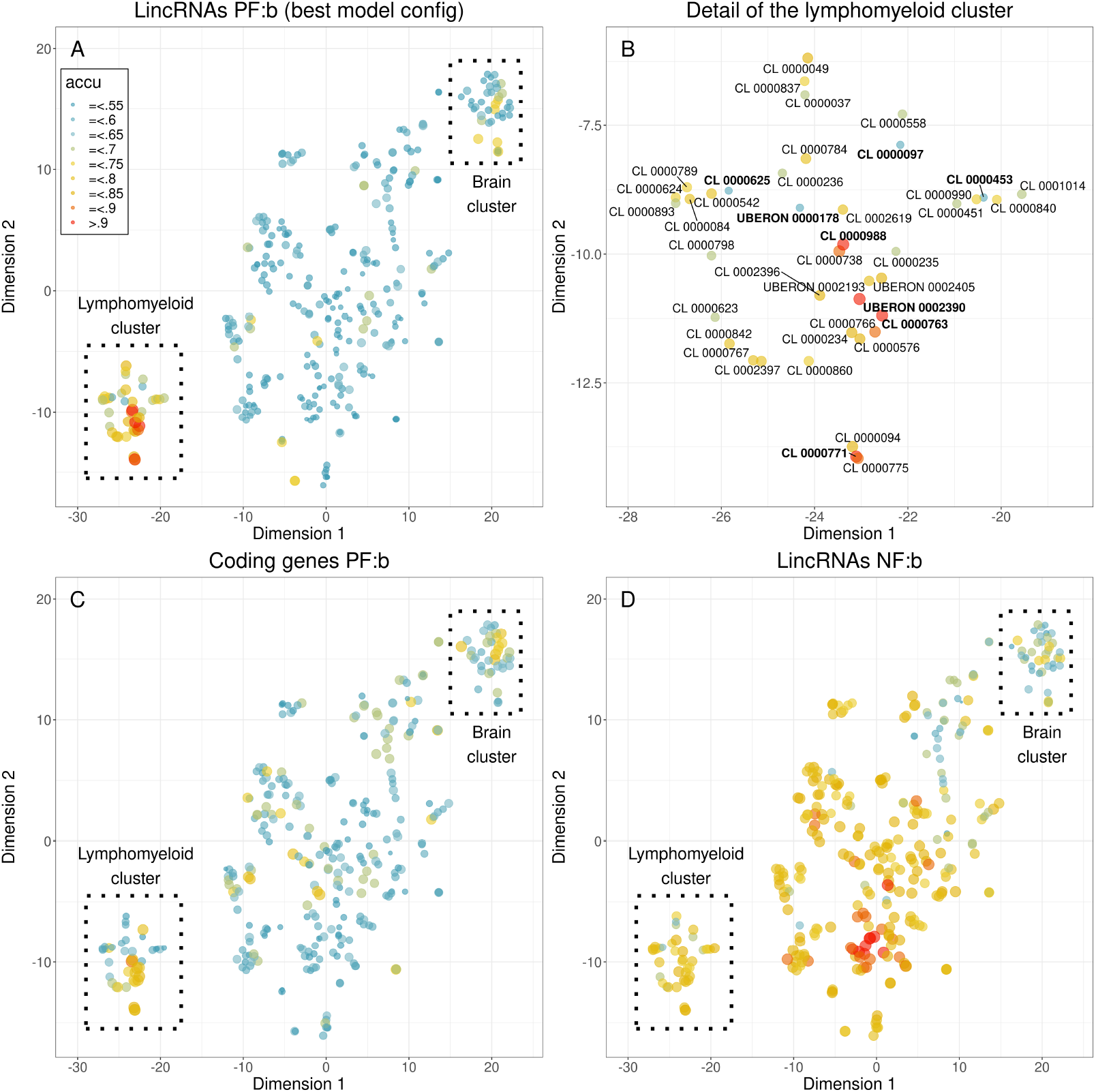
Visualisation of select model configurations across ontologies. Three model configurations were closely inspected across the FANTOM ontologies by mapping them using a *t*-SNE approach. Panels A and B: The best model configuration splits the map into two regions: on the one hand, the lymphomyeloid cluster with highly performing ontologies and, on the other hand, the remainder of the ontologies displaying low performances. The 4 most and 4 least performing ontologies of the lymphomyeloid cluster a highlighted in bold. Panel C: An identical model configuration but using coding genes instead of lincRNAs is mapped. Panel D: An identical model configuration but classifying NF:b instead instead of PF:b is mapped.

In the optimal model configuration, 21 out of 40 ontologies of the lymphomyeloid cluster achieved an accuracy of 75% or more and no other non-lymphomyeloid ontology reached that level of accuracy. Conversely, only 4 out of 40 ontologies of the lymphomyeloid cluster achieved an accuracy less than 65% compared to 282 (92%) of non-lymphomyeloid ontologies. The highest performance estimates, in the best configuration (cf. Fig. 7A), were observed for the hematopoietic system (UBERON 0002390), the hematopoietic cells (CL 0000988), the myeloid cells (CL 0000763) and the eosinophils (CL 0000771) with an accuracy of 93%, 92%, 91% and 91% respectively (see Fig. 7B). This is in line with the results obtained by [15] (on the very same dataset BH01) and by [17]. Among the lymphomyeloid cluster, the lowest performance estimates, in the best model configuration, were obtained for the following ontologies: CD8 positive alpha beta T cell (ontology CL 0000625), whole blood (UBERON 0000178), mast cell (CL 0000097) and Langerhans cells (CL 0000453) with an accuracy less than 60%. It must be noted that mast cells and Langerhans cells are immune cells not located in the bloodstream.

Again, in the optimal model configuration (Fig. 7A), the contrast in accuracy between the lymphomyeloid cluster and the non-lymphomyeloid cluster iwass the starkest. An identical model configuration except for the gene category in that it used coding genes achieved one of the best performance values too (Fig. 7C). However, the map allowed us to see that there was less contrast in terms of accuracy between the lymphomyeloid cluster and the non-lymphomyeloid cluster: the overall accuracy among the lymphomyeloid cluster was lower while, at the same time, relatively high-performing ontologies started to emerge throughout the non-lymphomyeloid cluster.

As observed earlier (Fig. 5), the average fragmentomic profile among the NF gene classes computed from the neutrophil ontology displayed a distinctive FD depression that set it apart from other gene classes. However, the model configurations involving the NF class appeared to achieve surprisingly low specificity and an accuracy map (Fig. 7D) allowed us to visualise the results across ontologies, highlighting the absence of contrast in accuracy when comparing lymphomyeloid and non lymphomyeloid ontologies.

## 3 Conclusion and discussion

This study focused on the capacity of a model to tell apart the tissue(s) of origin from other, decoy tissues based on fragmentation patterns observed in plasma-Seq data from healthy individuals and sought to identify the parameters most likely to improve the sensitivity and the specificity of the model. The tissue of origin was defined as a cluster of identifiable FANTOM ontologies related to lymphomyeloid cell types (in terms of expression landscapes).

The originality of our approach resided in assessing many model configurations to evaluate the tissue specificity conveyed by each parameter. Perhaps most notably, we looked at two gene categories: coding genes or long, intergenic, non-coding RNA genes (lincRNAs). We also looked at how tissue-specific gene classes could be defined to optimise the tissue identification. Finally, we looked at various combinations of fragmentomic features around TSSs.

In the optimal model configuration, good performance was achieved by most of the ontologies of the lymphomyeloid cluster (median accuracy greater than 75%). It can be reasonably assumed the low performance ontologies contribute no or little cfDNA to the bloodstream (because of physical barriers, low cell counts, low turn-over or a combination thereof). But high accuracy for a given ontology can either be indicative that the corresponding tissue or cell type may contribute a significant fraction of cfDNA or, another explanation, the result of functionally-similar yet different tissue or cell type contributing a significant amount of cfDNA.

It appears that lincRNAs using the tissue-specific genes (genes with a positive fold-change with most significant *p*-values in the FANTOM database) vs non-specific genes (as evidenced by high *p*-values) using all the features worked best (Fig. 4B).

Accuracy greater than 75% for most ontologies in the lymphomyeloid cluster (compared to 60% using coding genes). The impact of choosing a particular set of fragmentomic features was low in comparison to the choice of gene category and the method of building tissue-specific classes. By and large, the results can be explained by the average fragmentomic profiles of cfDNA in the vicinity of the genes classes (Fig. 5) and the high tissue-specificity of lincRNAs, documented in a number of studies [30–32]. This high tissue-specificity of lincRNA expression landscapes explains the superiority of lincRNAs in our study, particularly in terms of specificity, as the extent of the depression around a given TSS can only originate from a very limited set of identifiable cell types whereas the depression around coding genes’ TSSs can be expected to originate from a broader number of cell types, thus muddling tissue signatures. One obvious limitation of our study is that our parameters have been optimised for healthy samples. We believe that our results provide useful guidelines to test and validate future models in more specialised settings. In particular, since brain-specific lincRNAs are numerous, our results let us envision applications of this lincRNA-centred pipeline in the detection of the onset of neurodegenerative diseases [33].

Another novelty of this study is the use of tissue specificity (as provided by FANTOM) rather than raw expression levels in terms of fragments per kilobase million (FPKM) [14, 15, 34]. This raises interesting questions as to the means of quantifying tissue specificity as effectively as possible for liquid biopsy applications.

Our approach rests on the classification performance of a per-ontology gene classifier. Results can be partly, though not fully, interpreted in light of the average fragmentomic profiles around the TSSs of each gene class (see Fig. 5). It can be noted that the profile depression is often less pronounced in tissue-specific lincRNAs as compared to tissue-specific coding genes. This could be due to the lower transcription levels of lincRNAs [30–32, 35].

Both sorting methods (P sorting and FC sorting) worked similarly for coding genes. The situation was very different for lincRNAs, for which P sorting was vastly superior to FC sorting both in terms of sensitivity and specificity. We came up with a plausible explanation for the effectiveness of P sorting when it comes to lincRNAs is that lincRNAs are less intensely transcribed than coding genes and fold changes could be expected to be less precisely estimated for lincRNAs than for coding genes [30–32]. If true, *p*-values would offer an advantage over fold changes by providing a guarantee in terms of signal-to-noise ratio. However, this hypothesis was not supported by the FANTOM data. First, volcano plots representing fold-changes against *p*-values failed to support this explanation (data not shown). Also, the gene classes obtained by FC sorting and P sorting a no more similar in context of coding genes than they are in the context of lincRNAs (data not shown). The effectiveness of *p*-values in the identification of easy-to-classify gene classes thus remains to be explained.

Of course, other models integrating other descriptors, eg, epigenetic marks [36] or fragment sizes [17], other loci and based on a more diverse set of data must be designed to provide a tool usable at the bedside. We hope that our findings, in particular, those with regards to the use of noncoding RNAs, will help improve the sensitivity of ctDNA detection and help widespread liquid biopsy applications.

## 4 Acknowledgments

We’d like to thank Jennifer Dick (University of Sheffield) who helped revise the manuscript.

## 5 Data and code availability

Pre-processing and analysis were carried out by scripts written in Python (version 3.6.9) using NumPy (1.19.5), Pandas (1.1.5), Scikit-Learn (0.24.2) and Pysam (0.19.1). *t*-SNE was performed using R (4.2.2), Rtsne (0.15), data.table (1.14.0). The GRCh37 reference was used throughout this study, since that was the reference genome used for mapping the reads in the original study [15]. Scripts used to perform these steps are available.

BAM files are available from http://shendurelab.github.io/cfDNA/. The FANTOM databases are freely available from the FANTOM website: FANTOM DB1 from https://fantom.gsc.riken.jp/5/suppl/Hon_et_al_2016/data/assembly/lv3_robust/FANTOM_CAT.lv3_robust.info_table.gene.tsv.gz, FANTOM DB2 from https://fantom.gsc.riken.jp/5/suppl/Hon_et_al_2016/data/supp_table/supp_table_11.cell_type_gene_association.tsv, FANTOM DB3 from https://fantom.gsc.riken.jp/5/suppl/Hon_et_al_2016/data/assembly/lv3_robust/FANTOM_CAT.lv3_robust.CAGE_cluster.bed.gz, FANTOM DB4 from https://fantom.gsc.riken.jp/5/suppl/Hon_et_al_2016/data/expression/sample_ontology_association/ FANTOM DB5 is available as the supplementary table 4 in [24]. FANTOM DB6a from https://fantom.gsc.riken.jp/5/suppl/Hon_et_al_2016/data/expression/expression_atlas/FANTOM_CAT.expression_atlas.gene.lv3_robust.count.tsv.gz FANTOM DB6b from https://fantom.gsc.riken.jp/5/suppl/Hon_et_al_2016/data/supp_table/supp_table_10.sample_ontology_information.tsv, which were published alongside [23].Scripts used to perform these steps are available from our GitHub repository https://github.com/MuShuw/cnam_cfdna.

## Supporting information

S1. List of ontologies included in the lymphomyeloid cluster.

S2. A table containing summary statistics comparing lymphomyeloid and non-lymphomyeloid clusters is provided as supplementary data; the full list of results can be also be downloaded from the GitHub repository.

S3. Code to run a Bokeh interactive plot of the statistical descriptor of accuracy for all ontologies, lymphomyeloid ontologies and non-lymphomyeloid ontologies.

S4. An interactive HTML plot of the accuracy, across all ontologies, from the configuration PF:b using P sorting and the full feature set (FS11).

